# Formation and Gelation of Elastin-like Polypeptide Complex Coacervates

**DOI:** 10.1101/2025.01.15.633264

**Authors:** Rachel S. Fisher, Yihan Cheng, Lavinia Goessling, Allie C. Obermeyer

**Affiliations:** Department of Chemical Engineering, Columbia University, New York, NY 10027, USA

## Abstract

Protein liquid-liquid phase separation underlies the formation of membraneless organelles in cells and performs a key role in the assembly process of natural materials such as the assembly of tropoelastin into elastic fibers. Here, we engineered a series of charged elastin-like polypeptides (ELPs) that form complex coacervates, providing a rapid method to concentrate proteins into a fluid state. Compared to coacervates formed from simple coacervation, complex coacervates exhibited greater fluidity, likely due to differences between electrostatic interactions and hydrophobic forces. We designed these ELP’s to further contain crosslinking domains compatible with tyrosinase or transglutaminase and found that crosslinking was enhanced when proteins were in a complex coacervate compared to free in solution. Crosslinking the ELP complex coacervates led to the formation of gels with distinct properties dependent on the nature of the crosslinking. This work expands the design space of ELP hydrogels, offering a novel strategy for forming crosslinked networks from complex coacervates and providing opportunity for future use in tissue engineering and biocompatible biomaterials applications.

## Introduction

Protein-based materials present a promising option for sustainable materials, as they are naturally biocompatible and biodegradable, breaking down into non-toxic amino acids. Critically, nature offers a variety of biopolymers that match or even exceed the properties of synthetic polymers. However, until recently, their use was limited by production challenges. Advances in recombinant expression now allow for the production of these materials in relatively large quantities and enable easy modification of protein structures, tailoring biopolymer properties for specific needs.

Elastomeric proteins such as elastins, silks, and resilins, which possess a rubbery-like elasticity, are natural replacements for synthetic materials and have been used to make films, fibers, hydrogels and adhesives^1^. In nature, the properties of protein materials are more than the sum of their parts^2^. Elastomeric proteins assemble in such a way that they retain protein motion and flexibility required to stretch in response to force while forming enough cross-links to confer strength and stability. Recreating the carefully controlled biological processes that enable this hierarchical material assembly has long proven challenging. But, one mechanism common in the formation of many of these biomaterials is coacervation or phase separation (LLPS) ^2^. Notable examples of biopolymers that initially assemble through coacervation followed by crosslinking into fibers or gels include silks^3^, resilins, mussel foot proteins^4,5^, and tropoelastin^6^. The formation of a phase separated state as a precursor to self-assembly has several advantages. LLPS concentrates proteins providing a dense supply of precursor molecules; droplets can flow and coalesce allowing even coverage across molecular scaffolds; condensates can provide internal structure and molecular organization; and due to their rapid environmental response, small external changes can trigger changes in property or composition. With this in mind, we set out to design proteins capable of coacervation and crosslinking, mimicking a similar assembly process to that observed in nature.

Here we designed a series of recombinantly expressed elastin-like polypeptides (ELPs), containing cross-linking domains. These ELP’s are able to undergo complex coacervation, thereby increasing their concentration significantly, followed by enzymatic crosslinking to create a permanent network (Fig 1a). The material properties of condensates formed through LLPS are important in controlling their function [and to be useful in material production coacervates must conform to specific criteria]. By characterizing their viscoelastic properties as liquid coacervates and as crosslinked gels, we gain a deeper understanding of how protein sequence tunes material properties in both the liquid and gel phases.

**Figure 1.**
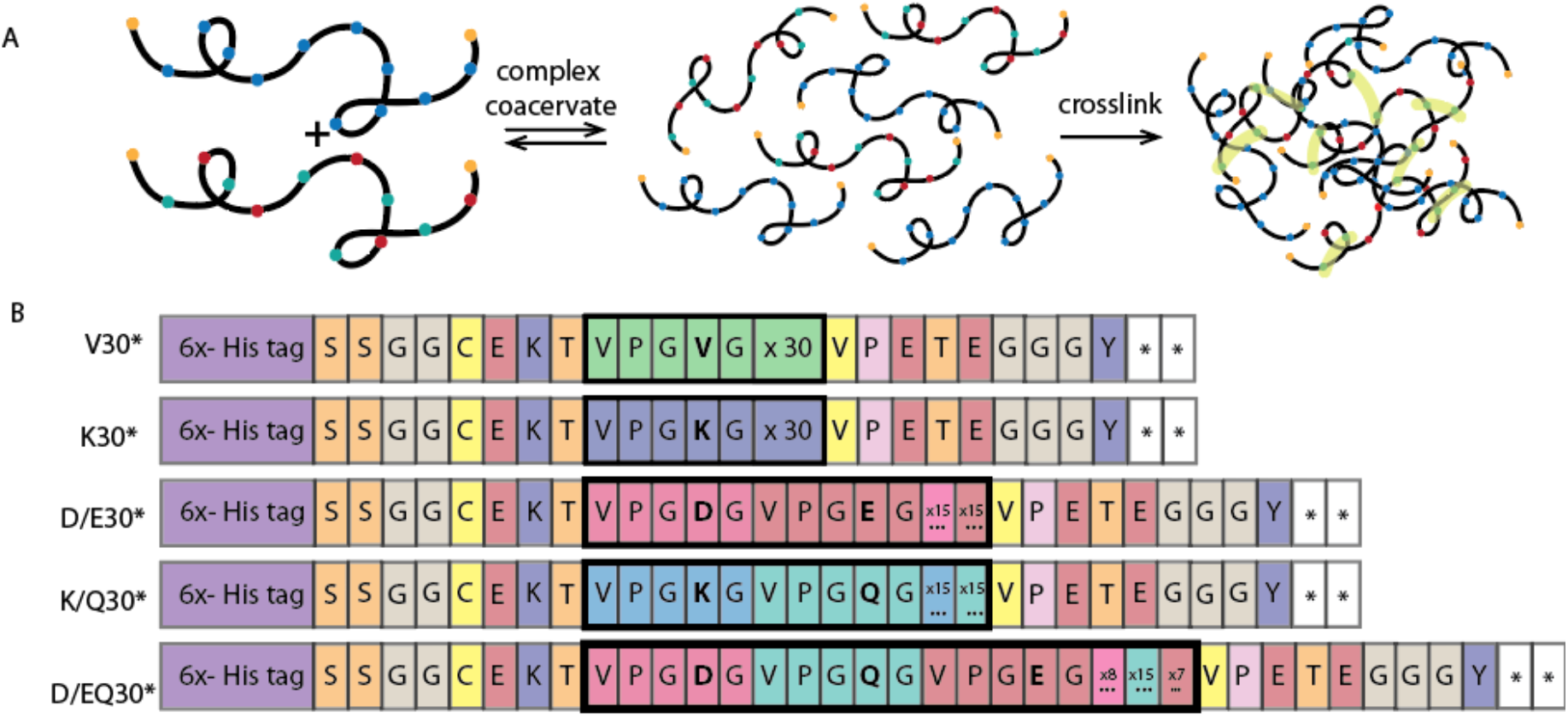
A) Schematic illustrating design principle: oppositely charged ELP’s, coacervation followed by crosslinking. B) Sequence of different elastin-like polypeptides used in this study.

## Results

### Design of ELPs capable of complex coacervation and crosslinking

With the aim of producing proteins that could be easily expressed, purified, concentrated via complex coacervation and then crosslinked, we designed a series of elastin-like polypeptides based on the well-studied VPGXG motif (Fig 1b). Naturally occurring elastins contain variations of a motif where X=V, and ELPs with this design are known to exhibit LCST behavior around room temperature, although this is dependent on length and salt concentration^7–9^. We therefore designed our initial ELP as 30 repeats of a V containing sequence (V30*). A His-tag was added at the C terminus to facilitate purification. To promote complex coacervation charged sequences were designed, the residue at position X was replaced with either a positive lysine residue (K30*) or negative glutamic or aspartic acid (D/E30*). While these designs would be capable of either simple or complex coacervation, the liquid-like droplets that form should dissolve upon changes in temperature or ionic strength, respectively. To create stable protein-based materials, enzymatic crosslinking sites were added at the C and N terminus of all sequences to allow end to end crosslinking by the enzyme tyrosinase (Fig 1a) ^10,11^. In addition to the potential for enzymatic chain extension, we also created designs capable of forming branched structures by the inclusion of internal enzymatic crosslinking sites. The addition of glutamine residues (Q) in half of the guest positions of K30* and D/E30* created KQ30* and D/EQ30* with crosslinking sites for transglutaminase, an enzyme that can form crosslinks between K and Q residues.

### ELPs undergo complex coacervation

We hypothesized that simple or complex coacervation of the ELP’s would provide a rapid and robust method to concentrate proteins to facilitate downstream material processing. We therefore assessed if the crosslinkable ELPs could undergo LLPS. The LCST of the neutral ELP (V30*) in 10 mM tris buffer without added salt was found to be approximately 43 °C, above which the protein formed simple coacervates (SI Fig 1). This transition temperature could be lowered to 33 °C with the addition of 500 mM NaCl and to below room temperature with the addition of 1.5 M NaCl (SI Fig 1).

Complex coacervates were formed by mixing ELPs containing a positive residue (K30* and KQ30*) with those containing a negative residue (D/E30* and D/EQ30*). By keeping the total protein concentration fixed and measuring turbidity as a function of mixing ratio we identified the ratio that results in the greatest amount of scattering, corresponding to the greatest amount of material. We found this occurred when there was a slight excess of positively charged ELP for all protein combinations (Fig 2 a). The formation of these condensates was highly sensitive to salt concentration and a complete loss of coacervation was observed upon addition of 50 mM NaCl when coacervates were composed of K30 and DE30 and 30 mM NaCl for coacervates containing an ELP with Q, and subsequently fewer charged residues (Fig 2 b). Optical microscopy was used to image samples at the peak turbidity ratio (Fig 2 c). We observed the formation of liquid-like droplets for all combinations. Interestingly droplets formed from K30 and DEQ30 appeared to have small aggregates inside them, an effect significantly more apparent when coacervates had been left to sit overnight (SI Fig 2). When we scaled the volume from around 100 µL to the mL scale we observed K30/DE30 and KQ30/DE30 form single semi-translucent phases that flow like a viscous liquid. K30/DEQ30 formed a dense, non-liquid two phase system we attribute to the majority of the sample forming individual aggregated complex (SI Fig 3 & 4). Consequently, we used only K30/DE30 and KQ30/DE30 in further experiments.

**Figure 2.**
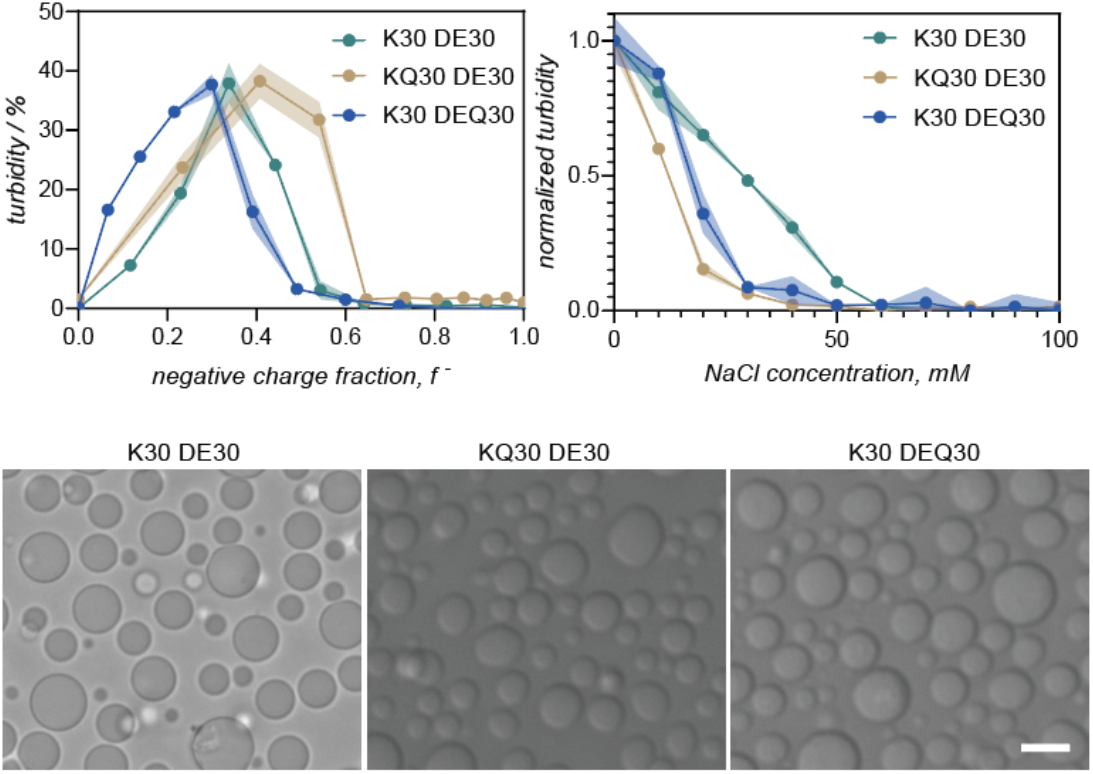
K30-DE30, KQ30-DE30, and K30-DEQ30 turbidity as a function A) charge fraction (f-) calculated as M^-^/(M^-^+M^+^) where M^-^ and M^+^ are the charge per negative and positive ELP. B) sodium chloride concentration C) Brightfield images of ELP coacervates at peak turbidity, 0 mM NaCl. Scale bar = 5 µm.

### Complex and simple coacervates have different viscoelastic properties

We next set out to determine the coacervate material properties. To investigate how sequence impacts viscoelastic response we performed DLS microrheology on simple coacervates of V30 and complex coacervates of K30/DE30 and K30/DEQ30 (Fig 3 a).

**Figure 3.**
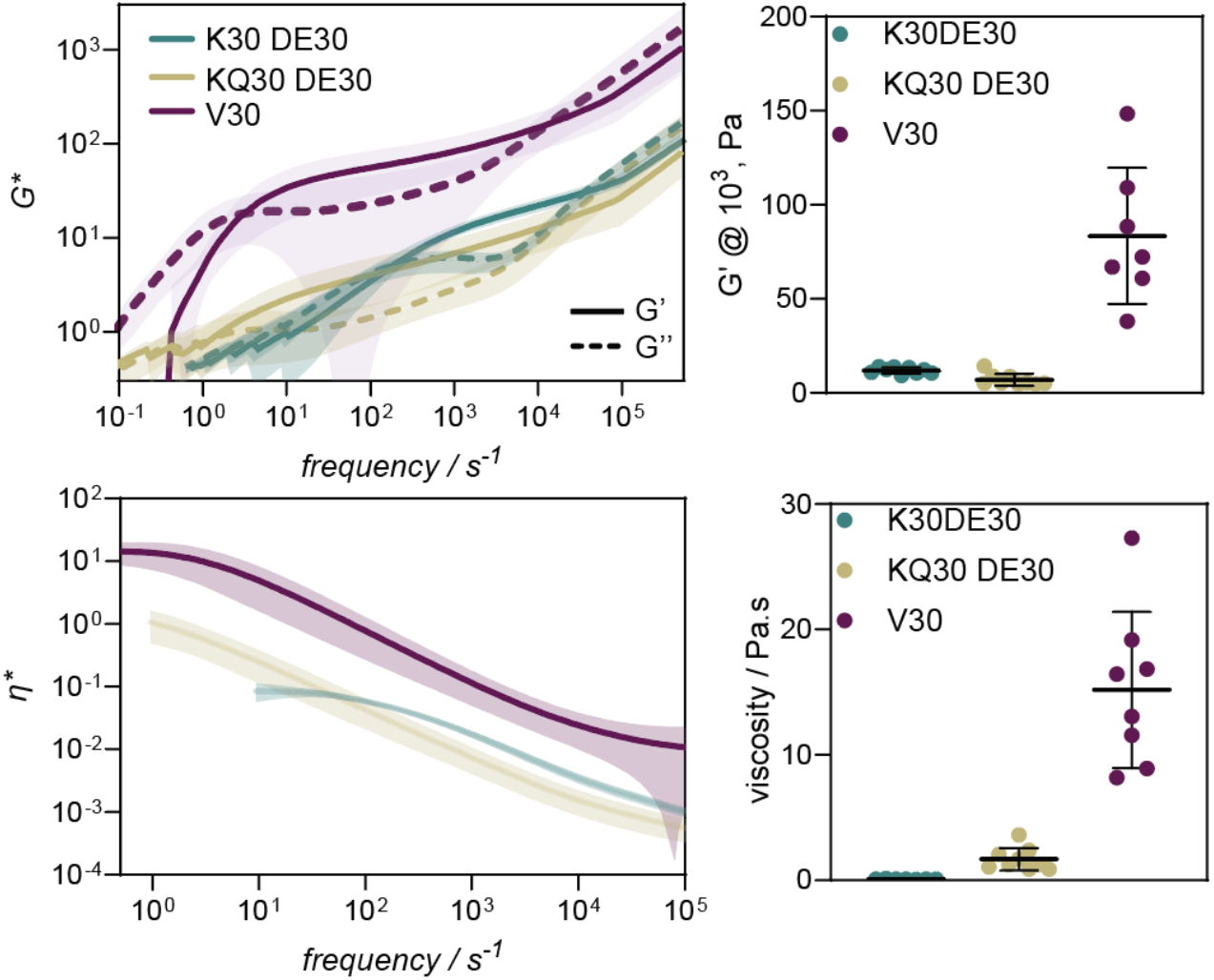
A) Complex modulus of K30-DE30 (10 mM tris, 37 C) (green), KQ30-DE30 (10 mM tris, 37 C) (gold) and V30 (10 mM tris, 500 mM NaCl, 37 C) ((purple). B) Modulus at 10^3^ s^-1^ C) Complex viscosity of K30-DE30 (green), KQ30-DE30 (gold) and V30 ((purple). D) Zero shear viscosity calculated from plateau region.

We were able to determine that both simple and complex coacervates were viscoelastic shear thinning fluids. We found a small decrease in elastic modulus between K30 and KQ30 condensates (Fig 3b), consistent with the lower charge of the KQ30 ELP. We also observed a longer crossover time, consistent with charged residues being further apart from each other (Fig 3c). Interestingly, we observed that while simple coacervates of V30 have a viscosity of 15 ± 6 Pa.s, a value similar to that measured for other ELP coacervates^12,13^, the viscosity of the complex coacervates are much lower: 1.7 ± 0.9 Pa.s for KQ30-DE30 and 0.1 ± 0.06 Pa.s for K30-DE30 (Fig 3). These results would also indicate a decrease in viscosity with increasing protein charge.

We attribute this difference in viscosity between simple and complex coacervates to different dominant interactions governing their behavior. The simple coacervation of ELP’s is known to be predominantly governed by hydrophobic domains and atomistic simulations show that the hydrophobic effect strongly contributes to the coacervate phase^14^. Based on the high sensitivity to salt concentration (Fig 2b) we attribute the dominant interactions in complex coacervates to electrostatics. It is well established that independently an increase in charge will lead to a more viscous coacervate but the interplay between charge and hydrophobicity is more complex. Recently Ramírez Marrero *et al* designed a co-polymer system where as they decreased either charge density or hydrophobicity they observed complex coacervate softening but as they systematically replaced charge domains with hydrophobic they first observed a softening of the material which they attributed to the decrease in charge density but at lower charge densities observed material stiffening^15^. This indicated that below a certain charge density hydrophobicity became more important. As proteins tend to have a significantly lower charge density than polymers it is likely this charge-hydrophobicity interplay will be of great effect. Our results indicate that the differing interactions dominating simple or complex coacervation does result in significantly different material properties for protein coacervates and the no charge, simple coacervate is stiffer than electrostatically dominated complex coacervates despite electrostatic interactions generally being considered a stronger interaction. As reentrant phase transition as a function of salt has been observed for many proteins^16^ and multiple proteins associated with intracellular phase separation can do so both alone or with RNA^17,18^, this observation is likely generally applicable to the material properties of many biomolecular condensates.

This difference in viscosity is also significant when considering processing coacervates into functional materials. To enable successful injection of salmine / sodium inositol hexaphosphate coacervates through a catheter into rabbit renal arteries, 1.2 M salt had to be added^19^. This reduced the coacervate viscosity from ∼ 40 Pa.s at physiological salt conditions to less than 1 Pa.s, at which point it became fluid enough to be suitable. Assuming similar viscosity requirements for ELP’s, simple coacervates would create too viscous a fluid to inject but complex coacervates would be within an appropriate range.

### Crosslinking complex coacervates produces gels with varying strength

We next aimed to assess the impact of crosslinking on the protein materials. We first looked at the effect of crosslinking with tyrosinase in dilute protein solution, which we expect to form end-to-end crosslinks between chains. We hypothesized that this chain extension would lead to a lower transition temperature for simple coacervation, as it is known that longer ELPs coacervate at lower temps^9^. However, we observed the opposite, an increase in transition temperature for V30 (SI Fig 5). We attribute this to cyclization of the protein, effectively decreasing the apparent molecular weight. This cyclic ELP is evident as a band with lower apparent molecular weight on a SDS PAGE gel (SI Fig 6). When tyrosinase is added to a 1 mg/mL solution of the charged ELP variants we see very few higher order species (dimers, trimers etc) forming for DE30. In contrast, for the cationic ELPs a modest increase in the formation of multimers was observed for K30, and significant fraction (70%) of crosslinked species were observed for KQ30 (SI Fig 7). This could suggest that the more charged ELPs are self-repulsive or do not interact well with the tyrosinase. To assess the impact of coacervation on crosslinking we added tyrosinase to samples containing oppositely charged ELPs at low ionic strength conditions. When tyrosinase is added to coacervate containing samples, an increase in the concentration of higher order species is observed indicating increased crosslinking in the high concentration coacervate phase compared to in solution.

To assess the effect of crosslinking occurring within the coacervate phase on material properties we performed microrheology. Looking at complex coacervates of K30/DE30 we observe that after addition of the crosslinking enzyme tyrosinase, there is a progressive increase in the crossover time in the viscoelastic spectra indicating bonds are lasting longer on average (Fig 4a). After ∼ 8 hours we no longer observe a crossover time indicative of a fluid, indicating bonding has become permanent and the coacervates have transitioned to a soft gel. Interestingly we do not observe any increase in elastic modulus, indicating bond strength and number remains largely the same. This suggests that despite crosslinks forming, the material properties remain dominated by the electrostatic interactions that govern the phase separation behavior. This is likely due to each protein containing only two tyrosinase crosslinking domains but 30 charged residues. The small decrease we observe in the modulus, we attribute to some fraction of material forming circular self-interacting structures.

**Figure 4.**
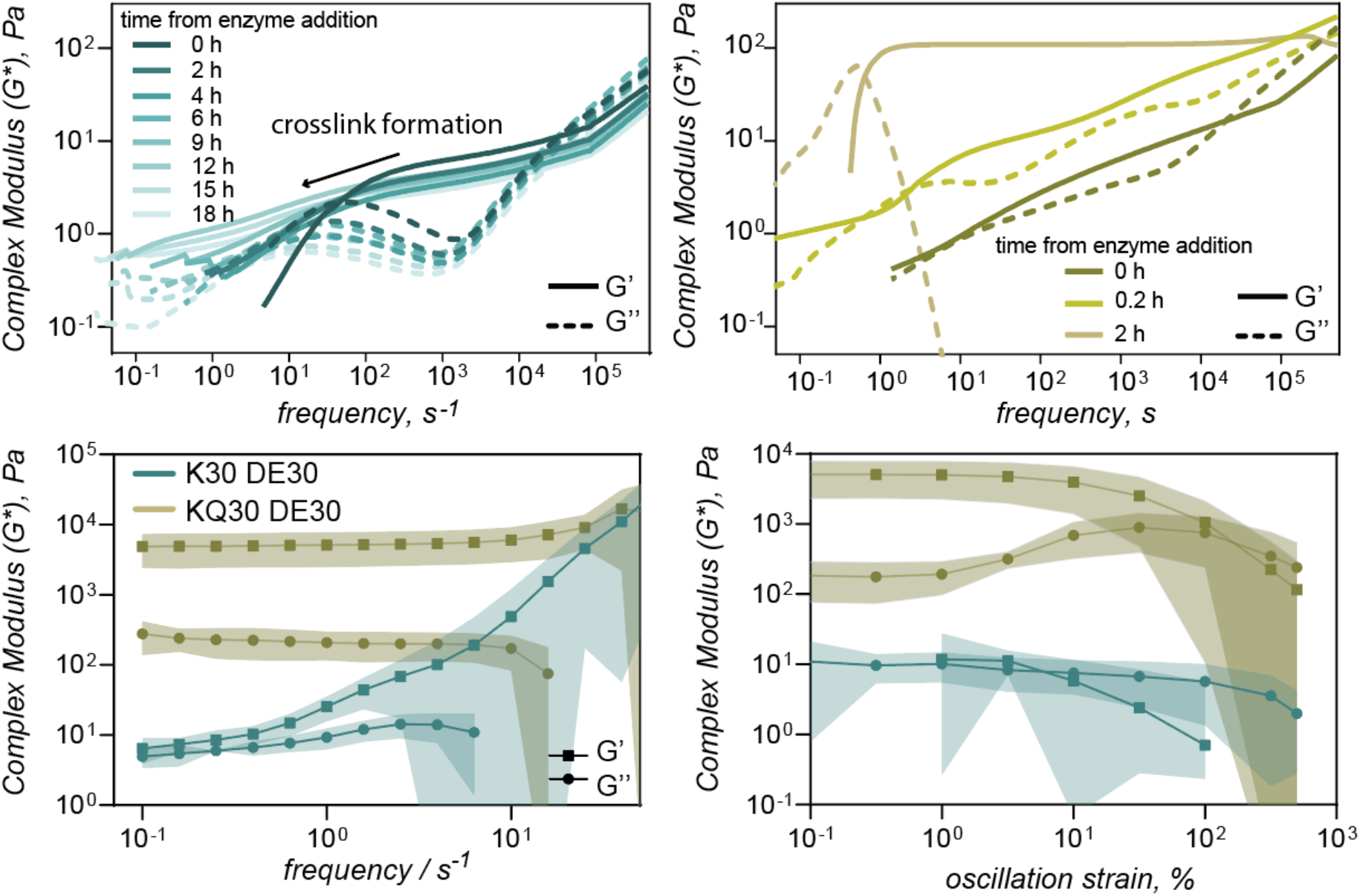
A) Complex moduli of K30DE30 at increasing times after addition of tyrosinase. B) Complex moduli of KQ30DE30 at increasing times after addition of transglutaminase. C) Storage and loss moduli of crosslinked coacervates of K30DE30 (green) and KQ30DE30 (gold) as function of the frequency. D) Storage and loss moduli of crosslinked coacervates of K30DE30 (green) and KQ30DE30 (gold) as function of the shear strain.

We next aimed to compare this chain extension crosslinking to those formed by transglutaminase between residues distributed along the length of the ELP. From analysis by gel electrophoresis, we observe that when ELP-KQ is crosslinked with transglutaminase in solution a similar fraction of higher order species from as when crosslinked with tyrosinase (SI Fig 6), However when crosslinking is performed in the presence of ELP-DE30 an increase in the formation of higher order species is observed (SI Fig 7) indicating enhanced crosslinking in the coacervate phase. From microrheology we observe that unlike end-to-end tyrosinase crosslinking, addition of transglutaminase appears to increase the modulus of the complex coacervates (Fig 4b). This indicates that more bonds, and in this case covalent bonds, are forming leading to increased material strength. The material rapidly becomes too stiff to reliably measure with via DLS microrheology.

To further investigate the microstructure of the crosslinked gels we characterized the viscoelasticity through low amplitude oscillatory shear measurements. We observed both K30-DE30 chain end crosslinked and KQ30-DE30 internally crosslinked materials behaved as gels over the entire frequency range studied, as indicated by a storage modulus (G’) greater than the loss modulus (G’’). However, there were substantial differences between the two gels: gels formed from KQ30 and DE30 had a storage modulus of ∼ 5±2 kPa, three orders of magnitude greater than that of the gel formed from K30 and DE30 with a storage modulus of ∼6±3 Pa (at 0.1 Hz). The KQ30-DE30 gel exhibits very little dependence of G’ on frequency, indicative of gel stability, comparatively K30-DE30 has a marked frequency dependence suggestive of a weaker gel with a low density of crosslinks^20^. This is consistent with our design with tyrosinase crosslinking creating a low density of crosslinks and subsequently a weak gel as compared to a stronger gel formed with the higher density crosslinks created with transglutaminase.

ELP’s alone or in tandem with other matrices have been used as tissue engineering scaffolds, where the modulus of the gel is a crucial element in controlling cell development. Human heart valve tissues range in storage modulus from 0.2-11 kPa^21^, and chemically cross-linked ELP hydrogels gels have demonstrated appropriate strength and utility as tissue scaffolds when in this range, indicating promise for the transglutaminase crosslinked gel developed here.

## Conclusion

We designed a series of ELPs that can form complex coacervates at low protein concentrations giving us a rapid technique to concentrate proteins into a fluid state suitable for injection or forming a thin film prior to crosslinking. We found significant differences in the viscoelasticity of ELP coacervates, with complex coacervation creating more fluid condensates than simple coacervation. We attribute this to hydrophobic interactions dominating simple coacervate behavior, whereas electrostatics dominate complex coacervation behavior. This finding is interesting when considering how different interactions impact the material properties of cellular condensates, demonstrating condensates governed by charged interactions may be more fluid. This is also instructive when designing ELP-based coacervates for processes involving injection, where a more fluid material may be preferential.

We found that while most of these proteins could be crosslinked in solution, the crosslinking yield increased under phase separating conditions and that this crosslinking in the coacervate phase led to network formation. Crosslinking of ELPs has been achieved using chemical agents^22–28^, resulting in gels with storage moduli in the range of 5-300 kPa with sequence, crosslinker, and crosslinker timing all influencing the hydrogel properties.

Crosslinking has also been using achieved in a more biocompatible fashion by photocrosslinking resulting in gels with storage modulus ∼1-3 kPa^29^ or enzymatically resulting in a gel with storage modulus of ∼0.2 kPa^30^. We compared two different enzymatic mechanisms of crosslinking, either through chain extension or via regular crosslinking along the biopolymer backbone, with potential crosslinking sites 5 residues apart residues. These produced two significantly different gels, one weak with a low crosslinking density and a more well-structured gel with a higher elastic modulus, with a similar order of magnitude as ELP hydrogels produced from chemical crosslinking. ELP-based materials have been extensively designed and shown great utility in tissue engineering applications, here we add a new biocompatible approach to the toolbox to create ELP hydrogels by creating crosslinked networks from ELP complex coacervates.

## Materials & Methods

### Cloning of ELPs

Five different ELP sequences were designed: D/E30, DE/Q30, K30, K/Q30, and V30, representing ELPs with anionic, cationic, or neutral residues in the guest position (X). All ELP sequences contained BbsI and BsaI restriction enzyme recognition sites at the N- and C-terminus, respectively, to allow for recursive directional ligation. Additionally, tyrosinase recognition sites (C at the N-terminus and EGGGY at the C-terminus) were added to enable protein-protein coupling for post-translational modification.

The valine containing ELP (V30) was originally obtained from Addgene (#67014). The V30 gene was transferred to a pETDuet plasmid by HiFi assembly. The type IIS restriction sites and the tyrosinase crosslinking sites were also added to the resulting plasmid via HiFi assembly.

The anionic ELP (D/E30), containing D and E in the guest position, was ELP-D/E30* in pET-24(+) plasmid was ordered from Twist.

The synthesis of the other ELP sequences involved polymerase chain cycling (PCA). Pairs of 100 bp-long forward and reverse primers were designed in such a way that each pair has a 30 bp-long overlapping region, with T_m_ at around 65 ºC, and no non-specific binding within five mismatches. Primers ordered from Integrated DNA Technologies were first annealed and then amplified through polymerase chain reaction (PCR). Restriction enzyme digestions using XhoI and XbaI were done on these PCR products and ELP-D/E30* in pET-24(+) plasmid so that the desired ELP sequences can be ligated into the same plasmid backbone. For ELP-DE/Q30* and ELP-K/Q30*, four pairs of long primers were used for PCA.

For synthesis of ELP-K30* to avoid non-specific bindings, shorter sequences (ELP-K10*) were first made using two pairs of long primers and then elongated via one or multiple recursive directional ligations. For recursive directional ligation, a double digestion with BbsI and BsaI was done on PCR amplified ELP sequences using T7 forward and reverse primers. A single digestion using BsaI followed by dephosphorylation using Quick CIP (NEB) was done on the full plasmid. The double digested ELP sequence and the single digested dephosphorylated plasmid were ligated using T4 ligase (NEB).

### Expression and Purification of ELPs

Plasmids with target ELP genes were first transformed into NiCo21 (DE3) *E. coli* host cells. For protein expression, a 5 mL overnight culture was inoculated in 1L LB media supplemented with 100 mg of kanamycin. After incubating at 37 ºC and shaking at 250 rpm for 24 hours without IPTG induction, cells were harvested by centrifugation and resuspended in either lysis buffer (300mM NaCl, 50mM NaH_2_PO_4_, pH 8.0) or 1x phosphate-buffered saline (PBS) solution, depending on the following purification method. The lysate was stored in -80 ºC and then thawed for sonication (2s on, 4s off cycle) with a total time of 30 minutes. Soluble proteins were separated from cell debris by centrifugation at 10,000 rpm for 45 minutes. For all cationic variants including K30* and K/Q30* both RNase A (225 µg/L of culture, from Qiagen) and DNase I (200 µg/L of culture, from Bovine Pancreas) were added prior to sonication, while polyethyleneimine (PEI) (1mL of 10% w/v PEI solution per 10mL of lysate, 764604 from Sigma-Aldrich) was added immediately after sonication to bind with and precipitate negatively charged impurities.

For V30* and D/E30 1x PBS solution was used as the resuspension buffer and inverse transition cycling (ITC) was used for purification. ITC takes advantage of the lower critical solution temperature (LCST) behaviors of ELPs. Based on the volume of lysate, Sodium chloride NaCl was added to a final concentration of 2M. For anionic ELP variants, after adding NaCl, solution pH was adjusted to 2.5 using 1M hydrochloric acid (HCl). The lysate was then heated and incubated in a 40 ºC water bath for 15 minutes. A hot spin was performed on the turbid lysate through centrifugation at 15,000 g for 15 minutes at 37 ºC. After carefully removing the hot spin supernatant, the ELP containing pellet was resuspended in 5∼10 mL of cold 1x PBS buffer and incubated on ice for 15 minutes before a cold spin by centrifugation at 15,000 g for 15 minutes at 4 ºC. The cold spin supernatant containing ELP was then separated from the solid contaminants.

For K30*, K/Q30*, and DE/Q30*, lysis buffer (300mM NaCl, 50mM NaH_2_PO_4_, pH 8.0) was used for resuspending harvested cells and FPLC (AKTA start) was used for purification. Multiple 5 mL HisTrap ™ HP columns were connected in series for FPLC. Lysis buffer with 35 mM or 250 mM imidazole was applied in the chromatography system as wash buffer and elution buffer.

Molecular weight and purity of target proteins were verified on SDS-PAGE before exchanging into 10 mM Tris buffer (pH 7.4) and spin concentration. For buffer exchange, 5mL HiTrap desalting columns (from Cytiva) connected in series were used in AKTA Start FPLC. Protein concentrations were quantified using Bicinchoninic Acid (BCA) assay with BSA (bovine serum albumin) as standard. All protein solutions were frozen in -20 ºC freezer for long term storage.

### Turbidity Measurements

Anionic ELPs or poly-L-aspartic acid sodium salt, x = 30 (PLD30 from Alamanda Polymers) was added to cationic ELPs or poly-L-lysine hydrochloride, x = 30 (PLKC30 from Alamanda Polymers) in Costar half area, flat bottom transparent polystyrene 96-well plates, and mixed by pipetting solutions several times. For all turbidity assays, the order of addition was kept the same (anionic > cationic). For both ELP and polymer solutions, concentrations were kept at 0.5 mg/mL. Right after mixing, absorbance at λ = 600 nm (A600) measurements were done with a Tecan Infinite M200 Pro plate reader. Percent transmittance (%T) was converted from triplicated A600 measurements.

For salt titration turbidity assay, various amount of concentrated NaCl solutions (1M or 5M) was first pipetted into 96-well plate. After water completely evaporated in 37 ºC incubator, cationic ELP or PLKC30 solutions was used to dissolve salt left in each well. Lastly, anionic component was added to induce coacervation. The mixing ratio used for salt titration turbidity assay was determined from the peak of turbidity assay.

### Optical Microscopy

Anionic ELPs were added to cationic ELPs in flat bottom transparent polystyrene 96-well plates coated with a 1% pluronics solution (PLF 127), and mixed by pipetting solutions several times. Imaging was performed on an EVOS™ FL Imaging System equipped with ×60/0.75 numerical aperture (NA) long-working distance Plan fluorite objective. ImageJ was used to further format and process images. Images were collected 1 h post coacervate formation.

### DLS Microrheology

DLS rheology measurements were performed as described by Cai et al^31^ and analyzed using their open access method (Version: 0.0.21) which can be found at https://dlsur.readthedocs.io/ and on GitHub at https://github.com/PamCai/DLSuR.

All measurements were performed in a Malvern low-volume quartz cuvette (ZEN2112). Coacervate samples (2 mL) were prepared by adding buffer, cationic ELP, 500 nm pegylated carboxy polystyrene beads, and anionic ELP in that order directly to the cuvette followed by pipetting to mix. The cuvette was then sealed with parafilm and left to stand for 10 min, before being placed into a falcon tube and centrifuged at (2000 xg) for 10 min to allow the condensed phase to settle to a layer at the bottom of the cuvette. Measurements were performed in this condensed phase without removal of the dilute phase so as to preserve equilibrium.

Autocorrelation curves were collected using a Malvern Zetasizer Nano ZS (633 nm laser) running version 7.12 Zetasizer software and operated in 173° non-invasive backscatter detection mode. The raw intensity autocorrelation function was collected at 4.2 mm, using automatic attenuator selection, for between 300 and 1200s. This was followed by fifteen 30 second measurements of the derived photon count rate at a series of positions, used to determine the scattering intensity. Data was exported and MSDs and complex moduli calculated as described in Cai et al^31^. Fitting data to extract cross over points and plateau moduli was performed in python.

### Rheological measurements

The rheological properties of the samples were characterized using the Discovery Hybrid Rheometer 30 (TA Instruments, USA). Samples were pre-gelled in 40 ml falcon tubes (SI Fig 4) and adjusted to diameter of 20 mm. A parallel plate of diameter 8 mm was used, and the geometry gap was set to 700 (±200) µm depending on sample thickness. Amplitude strain sweeps (0.1 – 500 %) were performed at a frequency of 1 Hz, frequency sweeps (0.1 – 100 Hz) at strain of 1 %.

## References

1 A. Miserez, J. Yu and P. Mohammadi, Chem. Rev., 2023, 123, 2049–2111.

2 A. Rising and M. J. Harrington, Chem. Rev., 2023, 123, 2155–2199.

3 A. D. Malay, T. Suzuki, T. Katashima, N. Kono, K. Arakawa and K. Numata, Sci Adv, 2020, 6, eabb6030.

4 E. Valois, R. Mirshafian and J. H. Waite, Sci Adv, 2020, 6, eaaz6486.

5 M. Renner-Rao, F. Jehle, T. Priemel, E. Duthoo, P. Fratzl, L. Bertinetti and M. J. Harrington, ACS Nano, 2022, 16, 20877–20890.

6 G. C. Yeo, F. W. Keeley and A. S. Weiss, Advances in Colloid and Interface Science, 2011, 167, 94–103.

7 D. W. Urry, T. L. Trapane and K. U. Prasad, Biopolymers, 1985, 24, 2345–2356.

8 D. W. Urry, Progress in Biophysics and Molecular Biology, 1992, 57, 23–57.

9 D. E. Meyer and A. Chilkoti, Biomacromolecules, 2004, 5, 846–851.

10 S. Halaouli, M. Asther, J.-C. Sigoillot, M. Hamdi and A. Lomascolo, J Appl Microbiol, 2006, 100, 219–232.

11 B. J. Kim, D. X. Oh, S. Kim, J. H. Seo, D. S. Hwang, A. Masic, D. K. Han and H. J. Cha, Biomacromolecules, 2014, 15, 1579–1585.

12 H. Betre, L. A. Setton, D. E. Meyer and A. Chilkoti, Biomacromolecules, 2002, 3, 910–916.

13 A. Vidal Ceballos, J. A. Díaz A, J. M. Preston, C. Vairamon, C. Shen, R. L. Koder and S. Elbaum-Garfinkle, Proceedings of the National Academy of Sciences, 2022, 119, e2202240119.

14 S. Rauscher and R. Pomès, eLife, 6, e26526.

15 A. Ramírez Marrero, L. Boudreau, W. Hu, R. Gutzler, N. Kaiser, B. von Vacano, R. Konradi and S. L. Perry, Macromolecules, 2024, 57, 4680–4694.

16 G. Krainer, T. J. Welsh, J. A. Joseph, J. R. Espinosa, S. Wittmann, E. de Csilléry, A. Sridhar, Z. Toprakcioglu, G. Gudiškytė, M. A. Czekalska, W. E. Arter, J. Guillén-Boixet, T. M. Franzmann, S. Qamar, P. S. George-Hyslop, A. A. Hyman, R. Collepardo-Guevara, S. Alberti and T. P. J. Knowles, Nat Commun, 2021, 12, 1085.

17 S. Maharana, J. Wang, D. K. Papadopoulos, D. Richter, A. Pozniakovsky, I. Poser, M. Bickle, S. Rizk, J. Guillén-Boixet, T. M. Franzmann, M. Jahnel, L. Marrone, Y.-T. Chang, J. Sterneckert, P. Tomancak, A. A. Hyman and S. Alberti, Science, 2018, 360, 918–921.

18 S. Elbaum-Garfinkle, Y. Kim, K. Szczepaniak, C. C.-H. Chen, C. R. Eckmann, S. Myong and C. P. Brangwynne, Proceedings of the National Academy of Sciences, 2015, 112, 7189–7194.

19 J. P. Jones, M. Sima, R. G. O’Hara and R. J. Stewart, Advanced Healthcare Materials, 2016, 5, 795–801.

20 T. G. Mezger, 2020.

21 T. Jiao, R. J. Clifton, G. L. Converse and R. A. Hopkins, Tissue Engineering Part A, 2012, 18, 423–431.

22 K. Di Zio and D. A. Tirrell, Macromolecules, 2003, 36, 1553–1558.

23 D. W. Lim, D. L. Nettles, L. A. Setton and A. Chilkoti, Biomacromolecules, 2007, 8, 1463–1470.

24 R. A. McMillan, K. L. Caran, R. P. Apkarian and V. P. Conticello, Macromolecules, 1999, 32, 9067–9070.

25 K. Trabbic-Carlson, L. A. Setton and A. Chilkoti, Biomacromolecules, 2003, 4, 572–580.

26 R. A. McMillan and V. P. Conticello, Macromolecules, 2000, 33, 4809–4821.

27 H. Wang, A. Paul, D. Nguyen, A. Enejder and S. C. Heilshorn, ACS Appl Mater Interfaces, 2018, 10, 21808–21815.

28 S. Vieth, C. M. Bellingham, F. W. Keeley, S. M. Hodge and D. Rousseau, Biopolymers, 2007, 85, 199–206.

29 Y.-N. Zhang, R. K. Avery, Q. Vallmajo-Martin, A. Assmann, A. Vegh, A. Memic, B. D. Olsen, N. Annabi and A. Khademhosseini, Advanced Functional Materials, 2015, 25, 4814–4826.

30 M. K. McHale, L. A. Setton and A. Chilkoti, Tissue Engineering, 2005, 11, 1768–1779.

31 P. C. Cai, B. A. Krajina, M. J. Kratochvil, L. Zou, A. Zhu, E. B. Burgener, P. L. Bollyky, C. E. Milla, M. J. Webber, A. J. Spakowitz and S. C. Heilshorn, Soft Matter, 2021, 17, 1929–1939.

