## Supporting Information for "Formation and Gelation of Elastin-like Polypeptide Complex Coacervates"

Elastin polypeptide sequence

| Protein | Sequence |
|---|---|
| V30* | MHHHHHHSSGGCEKTVPGVGVPGVGVPGVGVPGVGVPGVGVPGVGVPGVGVPGVGVP<br>GVGVPGVGVPGVGVPGVGVPGVGVPGVGVPGVGVPGVGVPGVGVPGVGVPGVGVPGVG<br>VPGVGVPGVGVPGVGVPGVGVPGVGVPGVGVPGVGVPGVGVPGVGVPGVGVPGVGVPG<br>VGVPGVGVPETEGGGYGWP* |
| D/E30* | MHHHHHHSSGGCEKTVPGEGVPGEGVPGDGVPGEGVPGDGVPGEGVPGDGVP<br>GEGVPGDGVPGEGVPGDGVPGEGVPGDGVPGEGVPGDGVPGEGVPGDGVPGE<br>GVPGDGVPGEGVPGDGVPGEGVPGDGVPGEGVPGDGVPGEGVPGDGVPGEGV<br>PGDGVPGEGVPETEGGGY* |
| D/EQ30* | MHHHHHHSSGGCEKTVPGDGVPGQGVPGEGVPGQGVPGDGVPGQGVPGEGV<br>PGQGVPGDGVPGQGVPGEGVPGQGVPGDGVPGQGVPGEGVPGQGVPGDGVP<br>GQGVPGEGVPGQGVPGDGVPGQGVPGEGVPGQGVPGDGVPGQGVPGEGVPG<br>QGVPGDGVPGQGVPETEGGGY* |
| K30* | MHHHHHHSSGGCEKTVPGKGVPGKGVPGKGVPGKGVPGKGVPGKGVPGKGVP<br>GKGVPGKGVPGKGVPGKGVPGKGVPGKGVPGKGVPGKGVPGKGVPGKGVPGKG<br>VPGKGVPGKGVPGKGVPGKGVPGKGVPGKGVPGKGVPGKGVPGKGVPGKGVPG<br>KGVPGKGVPETEGGGY* |
| K/Q30* | MHHHHHHSSGGCEKTVPGKGVPGQGVPGKGVPGQGVPGKGVPGQGVPGKGV<br>PGQGVPGKGVPGQGVPGKGVPGQGVPGKGVPGQGVPGKGVPGQGVPGKGVPG<br>QGVPGKGVPGQGVPGKGVPGQGVPGKGVPGQGVPGKGVPGQGVPGKGVPGQ<br>GVPGKGVPGQGVPETEGGGY* |

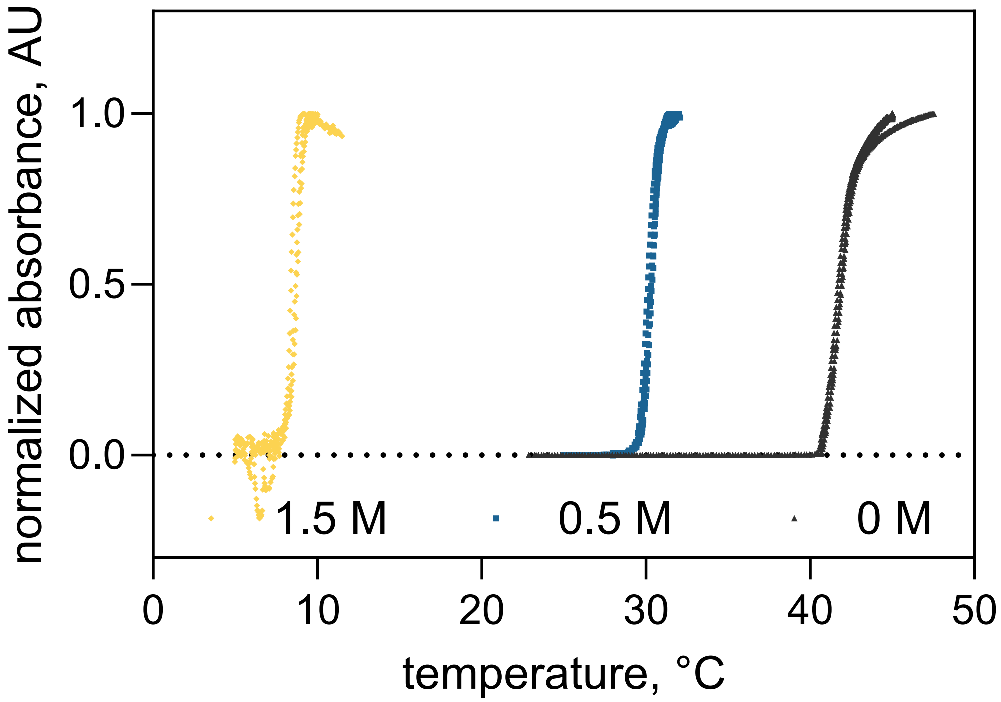


SI Fig 1) Normalized absorbance at 600 nm as a function of temperature at 0 M NaCl concentration (black), 0.5 M NaCl (blue) and 1.5 M NaCl (yellow) for samples of V30. Samples were prepared at 1 mg/mL of the protein in 10 mM tris buffer and heated at 0.3 °C/min to trigger the phase transition. Individual data points from three replicates are shown.

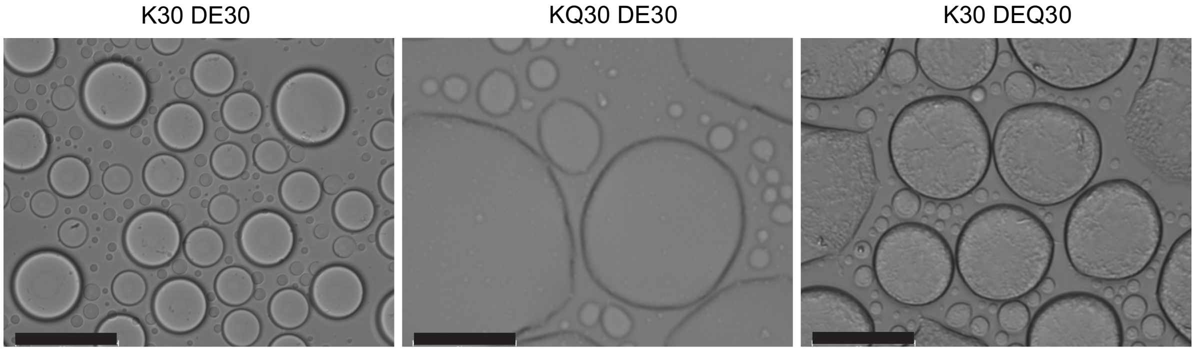

SI Fig 2. Brightfield images of ELP coacervates at peak turbidity, 0 mM NaCl after overnight incubation. Scale bar = 20 µm.

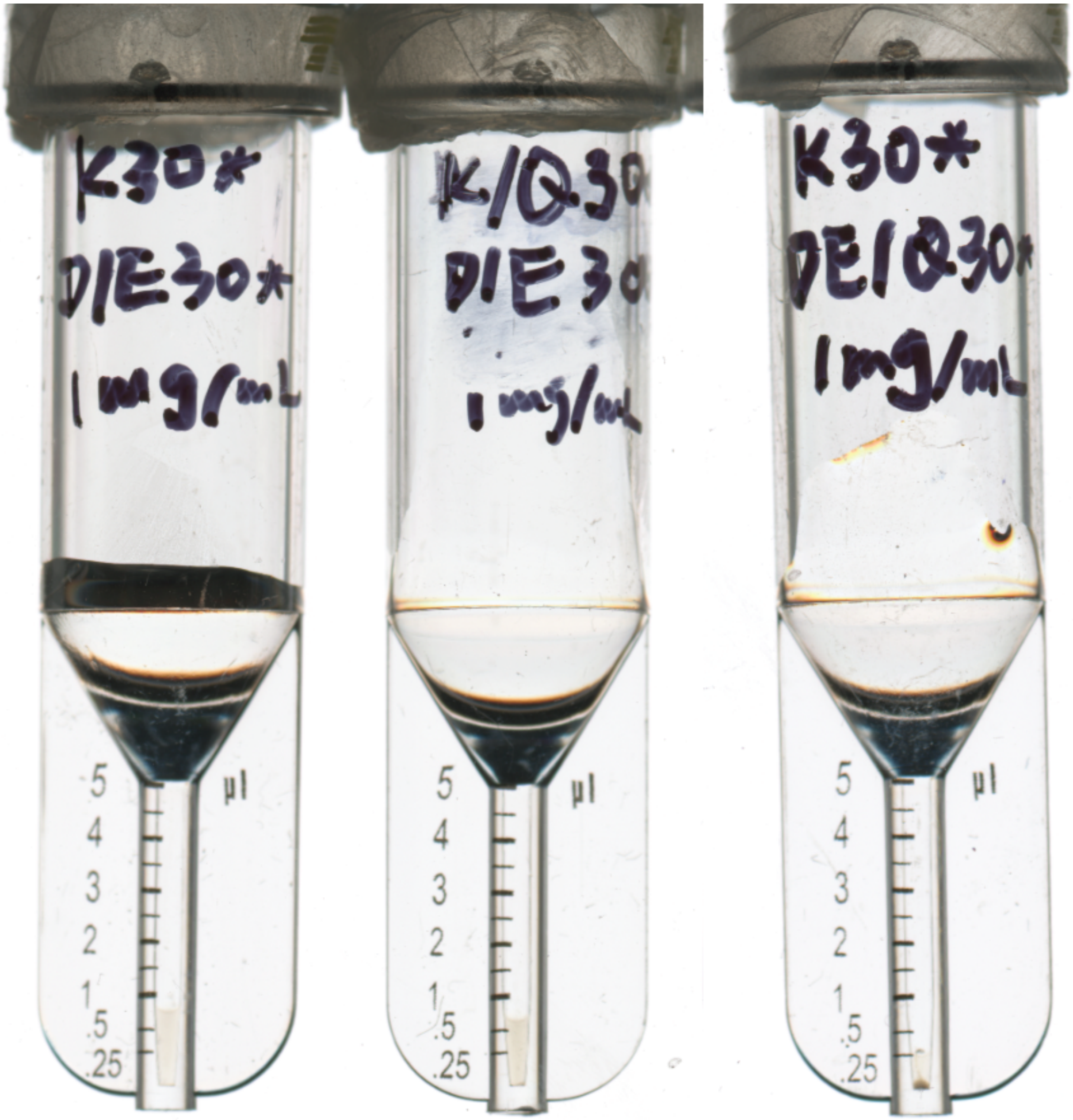


SI Fig 3. ELP coacervates were prepared with a mixture of charged ELPs at 1 mg/mL total protein concentration. Volume of the coacervate phase for K30+DE30 is approximately 0.7 µL from 100 µL total volume (left), and for KQ30+DE30 approximately 0.6 µL (middle). K30+DEQ30 forms a solid-like material in addition to a small volume (< 0.25 µL) of a liquid-like phase (right).

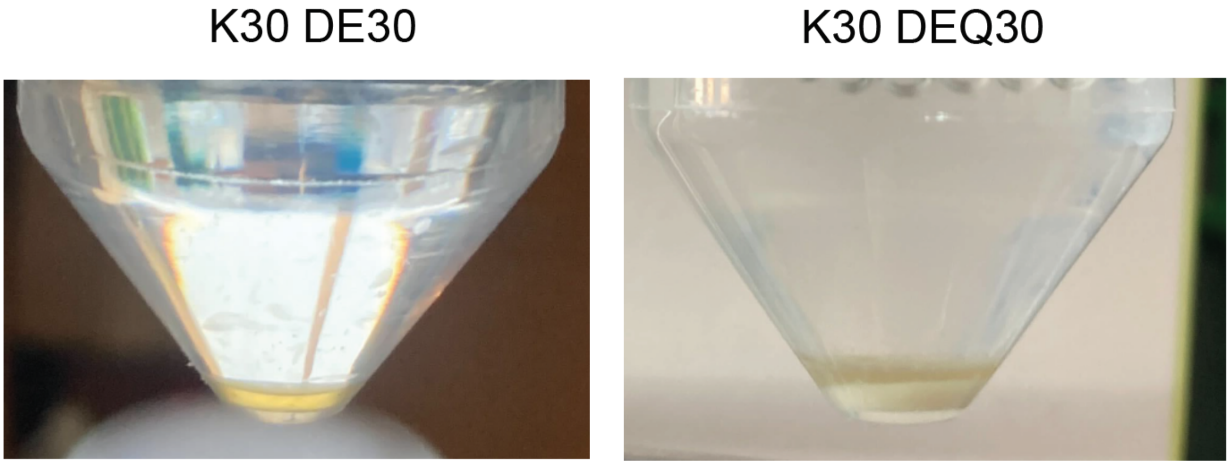


SI Fig 4. Coacervate phase formed from 20 mL of total volume for K30+DE30 (left) and from 40 mL of K30+DEQ30 (right). Coacervates formed with DEQ form a two-layer suspension.

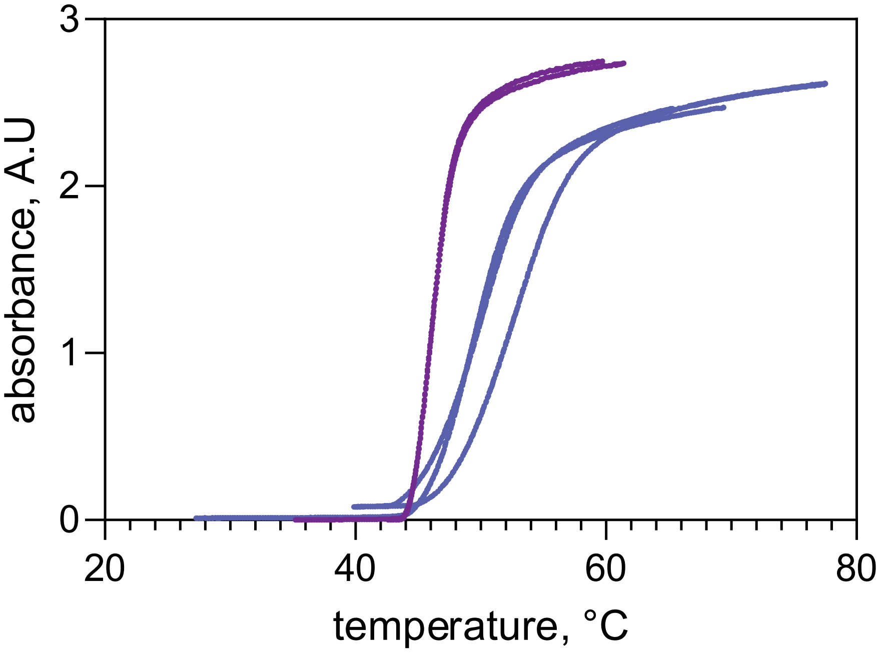


SI Fig 5) The turbidity of solutions of V30 (1 mg/mL) with tyrosinase (0.05 mg/mL) was monitored via absorbance at 600 nm as a function of temperature. Measurements were performed immediately after incubation with tyrosinase (0 h, purple) and after overnight incubation with tyrosinase (blue). Samples were heated at 0.3 °C/min to trigger the phase transition. Individual data points from three replicates are shown.

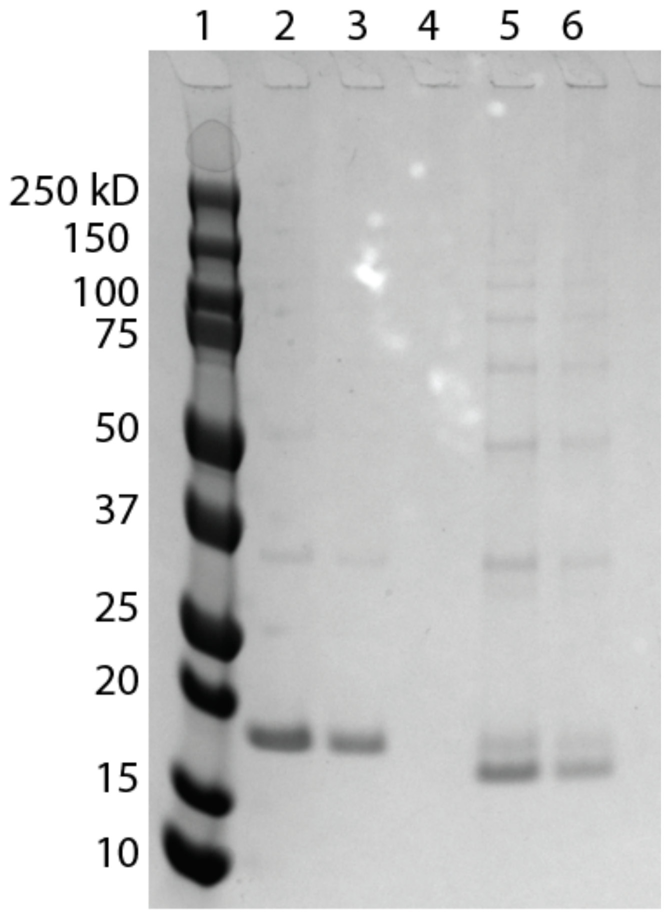


| | | | | | |
|---|---|---|---|---|---|
| tyrosinase | - | - | | + | + |
| % monomer | 91.5 | | | 17.3 | |
| % cyclic | 0 | | | 55.2 | |
| % HO species | 8.5 | | | 27.5 | |

SI Fig 6) SDS-PAGE gel analysis of tyrosinase mediated chain extension of V30. Lane 2 is V30 (0.25 mg/mL), lane 3 is V30 (0.125 mg/mL), lane 4 is tyrosinase (0.05 mg/mL), lane 5 is V30 (0.25 mg/mL) + tyrosinase (0.05 mg/mL) and lane 6 is V30 (0.125 mg/ml) + tyrosinase (0.05 mg/ml). The expected molecular weight of V30 is15.2 kDa and the band for pure V30 is observed at ~18 kDa. Following crosslinking with the enzyme additional high molecular weight bands are observed as well as a lower molecular weight band, which is proposed to be the cyclized polypeptide.

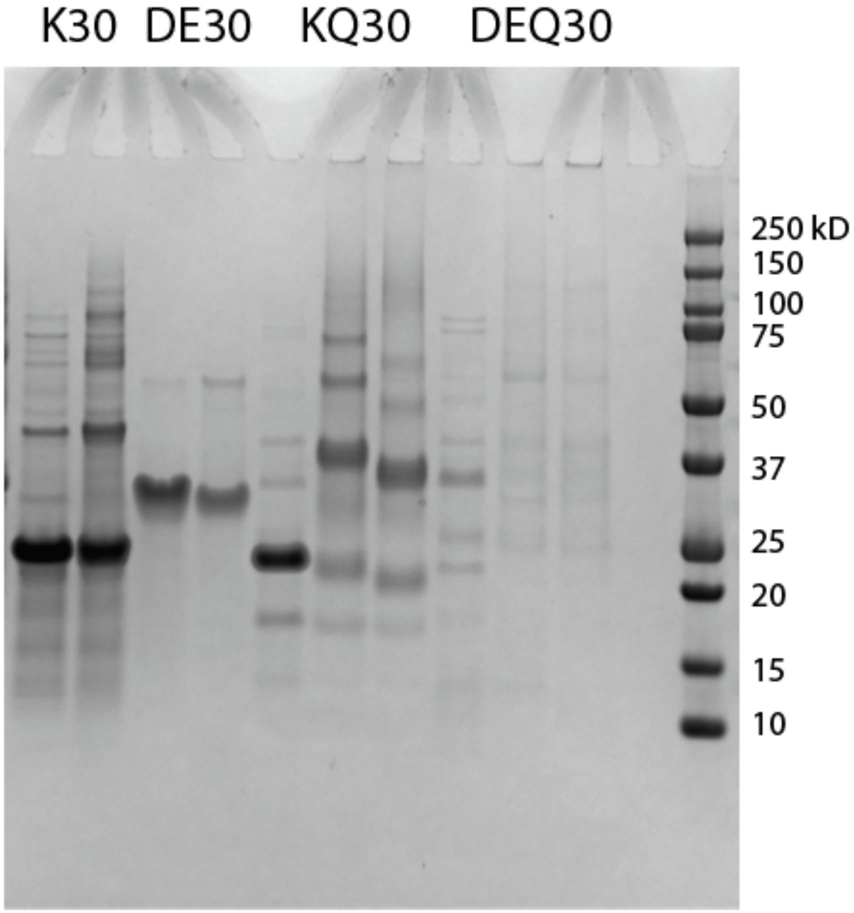


| ELP | K | K | DE | DE | KQ | KQ | KQ | DEQ | DEQ | DEQ |
|---|---|---|---|---|---|---|---|---|---|---|
| tyrosinase | - | + | - | + | - | + | - | - | + | - |
| transglutaminase | - | - | - | - | - | - | + | - | - | + |
| % monomer | 80 | 53 | 100 | 92 | 79 | 28 | 31 | 22 | | |
| % cyclic | 0 | 0 | 0 | 0 | 12 | 0 | 0 | 0 | | |
| % HO species | 20 | 47 | 0 | 8 | 9 | 72 | 69 | 78 | | |

SI Fig 7) SDS PAGE gel analysis of enzymatic crosslinking of ELP variants. Lane 1, K30 (1 mg/mL), lane 2 K30 (1 mg/mL) + tyrosinase (0.05 mg/mL), lane 3 DE30 (1 mg/mL), lane 4 DE30 (1 mg/mL) + tyrosinase (0.05 mg/mL), lane 4 KQ30 (1 mg/mL), lane 5 KQ30 (1 mg/mL) + tyrosinase (0.05 mg/mL), lane 6 KQ30 (1 mg/mL) + transglutaminase (0.05 mg/mL), lane 7

DEQ30 (1 mg/mL), lane 8 DEQ30 (1 mg/mL) + tyrosinase (0.05 mg/mL), lane 9 DEQ30 (1 mg/mL) + transglutaminase (0.05 mg/mL).

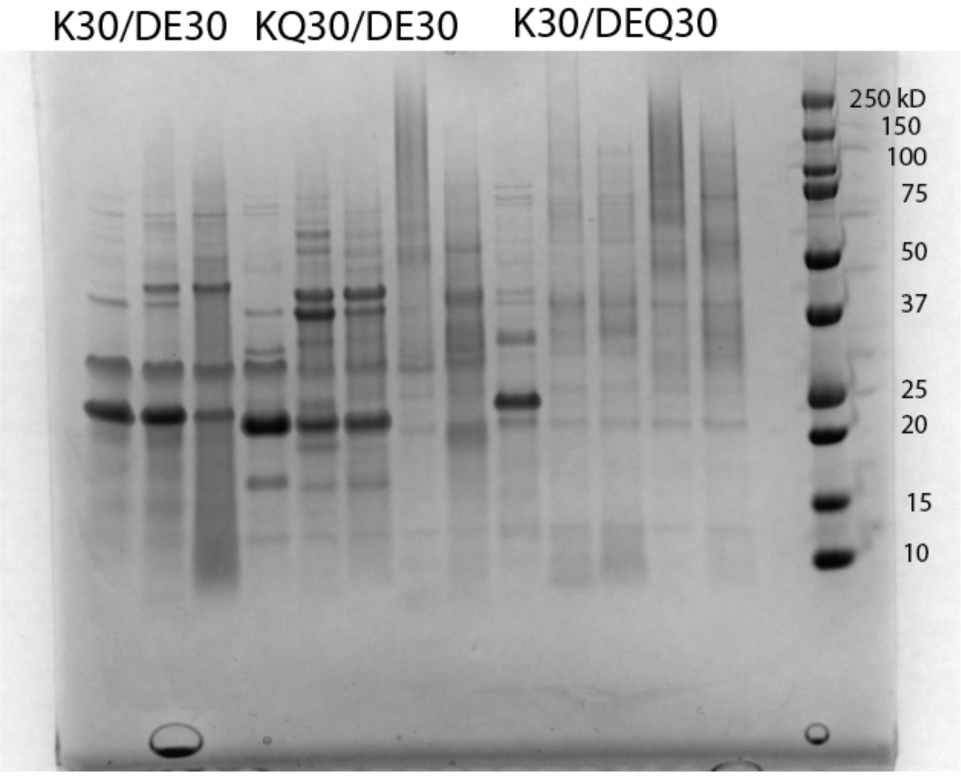


| | | | | | | | | | | | | | |
|---|---|---|---|---|---|---|---|---|---|---|---|---|---|
| +ive ELP | K | K | K | KQ | KQ | KQ | KQ | KQ | K | K | K | K | K |
| -ive ELP | DE | DE | DE | DE | DE | DE | DE | DE | DEQ | DEQ | DEQ | DEQ | DEQ |
| tyrosinase | - | + | + | - | + | + | - | - | - | + | + | - | - |
| transglutaminase | - | - | - | - | - | - | + | + | - | - | - | + | + |
| NaCl | - | - | + | - | - | + | - | + | - | - | + | - | + |
| % monomer | 83 | 62 | 60 | 72 | 33 | 51 | 18 | 42 | | | | | |
| % cyclic | 0 | 0 | 0 | 12 | 4 | 9 | 0 | 0 | | | | | |
| % HO species | 17 | 38 | 40 | 16 | 63 | 40 | 82 | 58 | | | | | |

SI Fig 8) SDS PAGE gel (top) annotated with band percentages (bottom). Lane 1 K30/DE30 (1 mg/mL, coacervates), lane 2 K30/DE30 (1 mg/ml, coacervates) with tyrosinase (0.05 mg/mL), lane3 K30/DE30 (1 mg/mL), 100 mM NaCl (prevents condensate formation) with tyrosinase (0.05 mg/mL). Lane 4 KQ30/DE30 (1 mg/mL, coacervates), lane 5 KQ30/DE30 (1 mg/mL, coacervates) with tyrosinase (0.05 mg/mL), lane 6 KQ30/DE30 (1 mg/mL), 100 mM

NaCl (prevents condensate formation) with tyrosinase (0.05 mg/mL), lane 7 KQ30/DE30 (1 mg/mL, coacervates) with transglutaminase (0.05 mg/mL), lane 8 KQ30/DE30 (1 mg/mL), 100 mM NaCl (prevents condensate formation) with transglutaminase, lane 9 K30/DEQ30 (1 mg/mL, coacervates) with tyrosinase (0.05 mg/mL), lane 10 K30/DEQ30 (1 mg/mL), 100 mM NaCl (prevents condensate formation) with tyrosinase (0.05 mg/mL), lane 11 K30/DEQ30 (1 mg/mL, coacervates) with transglutaminase (0.05 mg/mL), lane 12 K30/DEQ30 (1 mg/mL), 100 mM NaCl (prevents condensate formation) with transglutaminase.
